# Highly Dynamic Gene Family Evolution Suggests Changing Roles for *PON* Genes Within Metazoa

**DOI:** 10.1101/2022.05.17.492316

**Authors:** Sarah A.M. Lucas, Allie M Graham, Jason S Presnell, Nathan L Clark

## Abstract

Change in gene family size has been shown to facilitate adaptation to different selective pressures. This includes gene duplication to increase dosage or diversification of enzymatic substrates and gene deletion due to relaxed selection. We recently found that the *PON1* gene, an enzyme with arylesterase and lactonase activity, was lost repeatedly in different aquatic mammalian lineages, suggesting that the *PON* gene family is responsive to environmental change. We further investigated if these fluctuations in gene family size were restricted to mammals and approximately when this gene family was expanded within mammals. Using 112 metazoan protein models, we explored the evolutionary history of the *PON* family to characterize the dynamic evolution of this gene family. We found that there have been multiple, independent expansion events in tardigrades, cephalochordates, and echinoderms. In addition, there have been partial gene loss events in monotremes and sea cucumbers and what appears to be complete loss in arthropods, urochordates, platyhelminths, ctenophores, and placozoans. In addition, we show the mammalian expansion to three *PON* paralogs occurred in the ancestor of all mammals after the divergence of sauropsida but before the divergence of monotremes from therians. We also provide evidence of a novel PON expansion within the brushtail possum. In the face of repeated expansions and deletions in the context of changing environments, we suggest a range of selective pressures, including pathogen infection and mitigation of oxidative damage, are likely influencing the diversification of this dynamic gene family across metazoa.

## Introduction

The family of paraoxonase (*PON*) genes was named based on the discovery that one member could degrade the insecticide parathion, whose active metabolite paraoxon functions as a neurotoxic cholinesterase inhibitor(Costa et al. 1990). In 1996 it was revealed that this enzyme is part of a multigene family in humans(Primo-Parmo et al. 1996), whose genes were named *PON1, PON2*, and *PON3* in order of discovery. While these three genes produce protein products commonly known as serum paraoxonases, members of this protein family can also be found elsewhere. *PON1* and *PON3* are predominantly expressed extracellularly in the liver and their proteins are found on high-density lipoprotein (HDL) particles in blood serum. *PON2* is expressed intracellularly in a wide range of tissues and its protein product localizes to the endoplasmic reticulum and nuclear envelope(Ng et al. 2001; Horke et al. 2007). All three genes are in a tandem array on human chromosome 7, have approximately the same length, and contain the same number of exons. In contrast, non-mammal vertebrates like birds were revealed to only have a single *PON* gene. *PON*-like sequences have been identified in several other species including bacteria, nematodes, frogs, and mammals(Draganov and La Du 2004).

The native substrate(s) of PON proteins are still unclear. This makes it challenging to determine what selective pressure(s) resulted in the fixation and continued maintenance of these enzymes in most mammalian species(Billecke et al. 2000; Muthukrishnan et al. 2012). To address this, early physiological studies identified these proteins interact with a wide range of chemical structures. While PON1 hydrolyzed compounds such as paraoxon and lipid peroxides, the only classes of compounds which all three mammalian PONs can act upon are aromatic and aliphatic lactones (cyclic carboxylic esters)(Draganov et al. 2005; Bar-Rogovsky et al. 2013), arylesters (aromatic esters)(Billecke et al. 2000; Draganov et al. 2005; Khersonsky and Tawfik 2005), and homo-serine lactones which are key molecules for quorum sensing in bacteria(Draganov et al. 2005; Stoltz et al. 2007; Teiber et al. 2008). Other studies have revealed PON1 has atheroprotective - protection against plaque formation - effects(Shih et al. 1998; Tward et al. 2002), antioxidant properties through the degradation of lipoperoxides(Aviram et al. 1998), and an ability to co-regulate inflammation through an interaction with myeloperoxidase on HDL particles(Huang et al. 2013; Variji et al. 2019). Meanwhile PON2 has also been shown to have atheroprotective effects through its ability to reduce superoxide release(Ng et al. 2001; Horke et al. 2007; Altenhöfer et al. 2010; Devarajan et al. 2011) as well as anti-apoptotic properties(Krüger et al. 2015). PON3 has been associated with several diseases, but its exact functional role in those diseases have yet to be elucidated(Shih et al. 2007; Rull et al. 2012).

From this examination of human PON protein substrates and disease associations, it is highly suggestive this protein family plays two important roles: degrading lactones such as ones used in bacterial quorum sensing and contributing to antioxidant activity against lipoperoxides on HDL particles. However, there is evidence that in multiple independent lineages PON1 has been turned into a pseudogene, and that the loss of function may be adaptive, or the result of a relaxation of constraint in the aquatic environment(Meyer et al. 2018). An examination of the changes in *PON* copy number across metazoa and mammals could provide more clues as to what functions this enzyme provided.

The initial evolutionary study of this family found that *PON2* was the oldest member of this family followed by *PON3* and *PON1*(Draganov and La Du 2004); however, a more recent study challenged this finding. Through the incorporation of additional *PON* sequences, Bar-Rogovsky et. al determined that *PON3* diverged before *PON1* and *PON2*(Bar-Rogovsky et al. 2013). Since those papers were published, several marsupial and monotreme genomes have become available which would allow us a deeper look into the evolutionary history of this gene family in mammals to determine when it expanded in relation to the divergence of the different mammalian lineages. Additionally, both studies were limited in the number and types of genomes available to them. They were not able to investigate if these fluctuations in gene family size are restricted to mammals or how often it changed size in throughout metazoan evolution. With evidence of multiple independent expansions or deletions, we can begin to probe what functions are being selected for or against in this gene family.

Here, we explore the deep evolutionary history of the *PON* genes across 112 metazoan genomes and two choanozoan genomes. Ultimately, we determined that mammalian *PON* expansion occurred before the divergence of monotremata from the ancestor of all extant mammals (*i*.*e*., theria). In addition, we investigated the status of *PON* genes in a broad and diverse group of metazoans and found this mammalian expansion was not unique. Lastly, we highlight evidence of new specific duplications of *PON3* in the brushtail possum (*Trichosurus vulpecula*) that were followed by positively selected diversification. Overall, the contractions and expansions of the PON family suggest they are being acted upon by diverse evolutionary pressures such as combating bacterial biofilm formation or managing oxidative stress.

## Methods

### Identification of Metazoan *PON* family members

The 101 species used in this analysis (see Supplementary Table 1) were sampled by multiple research groups under variable conditions, and hence likely vary in the completeness of their gene content. Our approach to minimize false negative findings (i.e., false losses) was to examine multiple species within each group when possible, and to conservatively claim loss of a gene only when it was not detected in all species examined within the respective group.

*PON* genes were identified using a combination of HMMR searches and phylogenetic verification. The *PON* family of genes are characterized by the presence of an arylesterase domain(Primo-Parmo et al. 1996; Rodrigo et al. 1997); thus we used *hmmsearch* in HMMR 3.0(Eddy 1998) to search proteins for motifs that matched PFAM profile for arylesterase (PF01731.21) – this model was created using 1047 known *PON* sequences from 511 species across Eukaryotes. *PON* sequences were identified based on maximum full sequence e-value and maximum the best domain e-value of 1e-6 regardless of the number of domains present. If multiple *PON*s were identified within the same species, *PON*s identified on different chromosomes or on the same chromosome/scaffold, but > 100kb away, are designated by a different alphabet character. *PON*s on the same chromosome/scaffold and within 100kb are given the same alphabet character and a different number.

We then used PASTA (v.1.8.6)(Mirarab et al. 2015) to generate a multiple sequence alignment using default parameters except --mask-gappy-sites=6. The alignment was trimmed using Clipkit’s smart-gap mode and default parameters(Steenwyk et al. 2020). RAxML generated the optimal phylogenetic tree from twenty random starting trees and using the protgammaauto option. Then using the Le-Gascuel (LG)(Le and Gascuel 2008)+G amino acid substitution model, 1000 bootstraps were performed. Trees were visualized using FigTree (version 1.4.4, http://tree.bio.ed.ac.uk/software/figtree/) and modified in Adobe Illustrator (2020). The ambulacraria tree was produced using the same sequences identified and methods used for the metazoan analysis with the exception that only 200 bootstraps were performed.

### Identification of Mammalian *PON* family members

To identify mammalian *PON* family members, sequences (see Supplementary Table 2) were queried against human PON1 (NP_000437.3), PON2 (NP_000296.2), and PON3 (NP_000931.1) using BLASTP(Altschul et al. 1990) with a minimum query coverage of 90% and minimum percent identity of 50%. Additional criteria for identification were that the placental mammals had to have at least one of their PON proteins curated in RefSeq, and each protein must come from a unique chromosomal locus. In the case where multiple isoforms were available, the longest sequence was used.

Sequences identified using the criteria above were then aligned using webPRANK (https://www.ebi.ac.uk/goldman-srv/webprank/)(Löytynoja and Goldman 2010), and regions of high divergence or single species specific indels were trimmed manually using AliView (version 1.27)(Larsson 2014). Phylogenetic analysis of this alignment was done using PhyML (version 3.3.20190909, http://www.atgc-montpellier.fr/phyml/)(Guindon et al. 2010) using SPR tree improvement and 3 random starting trees with 200 bootstraps. Smart Model selection (SMS)(Lefort et al. 2017) determined the Jones-Taylor-Thornton (JTT) substitution model(Jones et al. 1992) with a gamma distribution (G) (parameter = 1.29) and 0.068 proportion of sites being invariable (I) was the best model using Akaike information criterion (AIC)(Akaike 1974). Trees were visualized using FigTree and modified in Adobe Illustrator (2020).

### Model Comparison: *PON* family duplication

To determine which of the three mammalian *PON*s was the first to diverge, three different tree models were generated based upon the tetrapod species tree. We then tested which of the three models best fit the tetrapod multiple sequence alignment using the CODEML program in PAML (version 4.9)(Z. Yang 2007). The JTT substitution model was used. The log-likelihood scores produced by CODEML were compared to determine statistical significance using a likelihood ratio test and a chi-square distribution with one degree of freedom(Huelsenbeck and Crandall 1997).

To confirm that the monotreme *PON* was not the result of merging of two *PON* genes, multiple sequence alignments of the individual exons and grouped exons as determined by NCBI gene were generated using PRANK. To determine where the monotreme sequence clustered, PhyML was used to generate a maximum likelihood tree (tree improvement: SPR, Number of random starting tree: 3, perform bootstrap: 200). Different nucleotide substitution models were used depending on what SMS determined.

### RNA-seq mapping

To verify that the *PON* expansion in the brushtail possum *(Trichosurus vulpecula)* is real we investigate whether RNA-seq samples could map to that region. Samples (see Supplementary Table 3) were downloaded from NCBI-SRA using sra-toolkit (v2.10.0, https://github.com/ncbi/sra-tools)(Anon). Reads were trimmed using trim-galore (0.4.4, cutadapt v1.14 https://github.com/FelixKrueger/TrimGalore)(Martin 2011; Krueger 2012), and had quality assessment done by fastqc (0.11.4), https://www.bioinformatics.babraham.ac.uk/projects/fastqc/)(Andrews 2010). Using BWA-MEM (v. 2020_03_19) (Van der Auwera et al. 2013), the reads were mapped to manually concatenated *PON* mRNA (XM_036759253.1, XM_036759536.1, XM_036759540.1, and XM_036761169.1) and visualized by Integrated Genome Viewer (IGV)(Thorvaldsdóttir et al. 2013) (v2.9.2).

### Testing for Positive Selection

To determine if the brushtail PON3, PON4A, and PON4B proteins were experiencing positive selection, RefSeq mRNA, excluding the stop codons and untranslated regions, of the marsupial PON3 proteins were acquired from NCBI Nucleotide. The nucleotide sequences were aligned using PRANK and spurious sequences, UTRs, and stop codons were manually trimmed. We used a likelihood-ratio test to compare the M1 and M2 models and M7 and M8 models within the CODEML package in PAML to determine if positive selection was detected across all branches (options Model = 0, NSsites = 1 2 7 8)(Yang and Swanson 2002; Ziheng Yang 2007). We also tested if positive selection was occurring on specific branches involving the duplication within the brushtail PON proteins using PAML’s branch-site model test 2 using the same multiple sequence alignment from the previous positive selection tests (options. Model = 2, NSsites = 2, fix_kappa = 0, kappa = 2, omega = 1, fix_alpha = 1, alpha = 0)(Zhang et al. 2005). All sites which were identified by the Bayes Empirical Bayes analysis were taken to be under positive selection(Yang et al. 2005). Additionally, another branch-site analysis was also done using BUSTED (https://www.datamonkey.org/analyses)(Murrell et al. 2015) using the same alignment as the PAML analysis. The three branches leading toward the brushtail *PON* sequences as well as the internal node connecting *PON3* and *PON4B* were selected as being in the foreground.

### Protein Modeling

To model where the predicted positively selected sites were located, an amino acid multiple sequence alignment between the marsupial and rabbit PON3 proteins was generated using PRANK(Löytynoja and Goldman 2010). Positively selected sites were then mapped onto the rabbit serum paraoxonase (protein data bank: 1V04)(Harel et al. 2004). The protein was visualized using Chimera 1.13.1(Pettersen et al. 2004). Visually, it appeared that the sites experiencing positive selection appeared to be clustered near the active site of PON1. To statistically confirm if this was indeed the case, we used GETAREA to identify solvent exposed surface residues in the crystal structure(Fraczkiewicz and Braun 1998) and used that information to run random permutation simulations to statistically determine if these residues are clustered(Clark et al. 2007).

## Results

### *PON* expansion and contraction has occurred multiple independent times within metazoa

HMMER was used to identify the sequences containing an arylesterase domain. It is the only functional domain found within *PON*s and is unique to this protein family. Across 99 metazoan species and two choanozoa, 41 species did not contain a protein with a high confidence arylesterase domain, including species within ctenophora, placozoa, platyhelminth, urochordata, or arthropoda; however, the remaining 60 species within porifera, cnidaria, chordata, ambulacraria, spiralia and ecdysozoa showed broad evidence of *PON* genes. All 159 sequences were aligned and subjected to phylogenetic analysis.

The resulting phylogeny provides evidence of ancestral duplication among closely related species (Fig 1, Fig S1). Within echinoderms, tardigrades, and bivalves, there is evidence of a single ancestral duplications which resulted in two ancestral PON genes for each taxonomical group. Similarly, there is also evidence of two ancestral duplications in cephalochordates and mammals, resulting in three ancestral PON genes.

**Figure 1.**
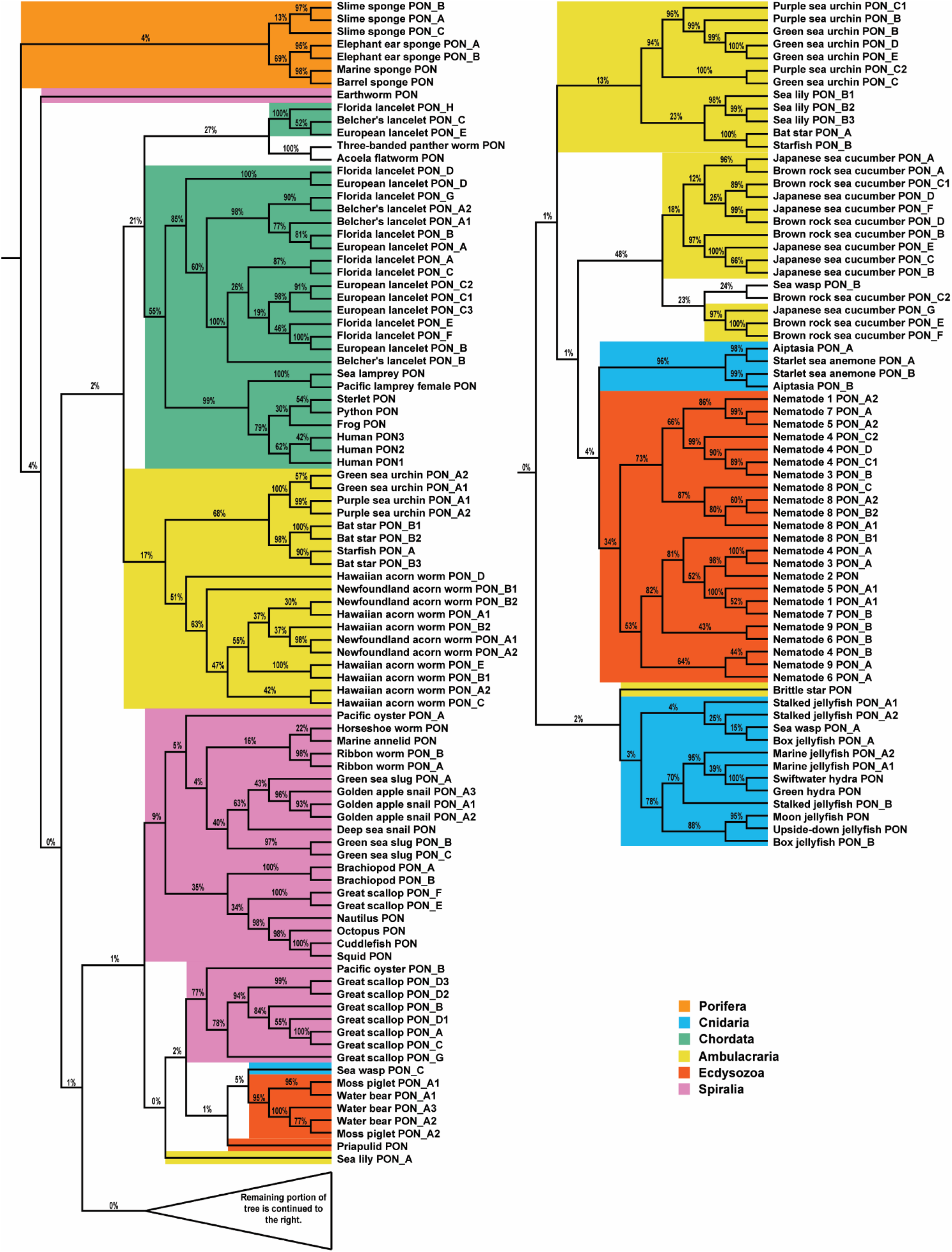
Evolution of PON in metazoans. Phylogenetic tree of PON family proteins in metazoans determined by RAxML based on multiple sequence alignment. Bootstrap support values are shown as percentages out of 1000 bootstraps. If species have multiple PON genes and they are located on different chromosome/scaffold or are sufficiently far from one another, then they are indicated by different alphabet characters. If they PON genes are located on the same chromosome/scaffold and are within one 100 kb of another PON gene, this is indicated by a number.

In addition to those instances of ancestral duplications, there are many instances of lineage specific PON expansions through tandem duplication and other gene duplication mechanisms. Two examples of tandem duplication include *PONA1* & *PONA2* for the green sea urchin (*Lytechinus variegatus*) and purple sea urchin (*Strongylocentrotus purpuratus)*. Both species had an independent tandem expansion of the same PON gene as the sequences within a species cluster better together than with a *PON* from the other species and are located on the same chromosome. This contrasts with the ancestral tandem duplication observed in mammals (Fig 2A). Besides tandem duplication, there is evidence of additional gene duplication, particularly in the Great Scallop (*Pecten Maximus)*. While three of its nine *PON* genes are in a tandem array (i.e. less than 100 kb away from each other) on one of its chromosomes, the remaining six are scattered among five chromosomes and scaffolds (Supplementary Table 1) giving evidence to some alternate gene duplication mechanisms.

**Figure 2.**
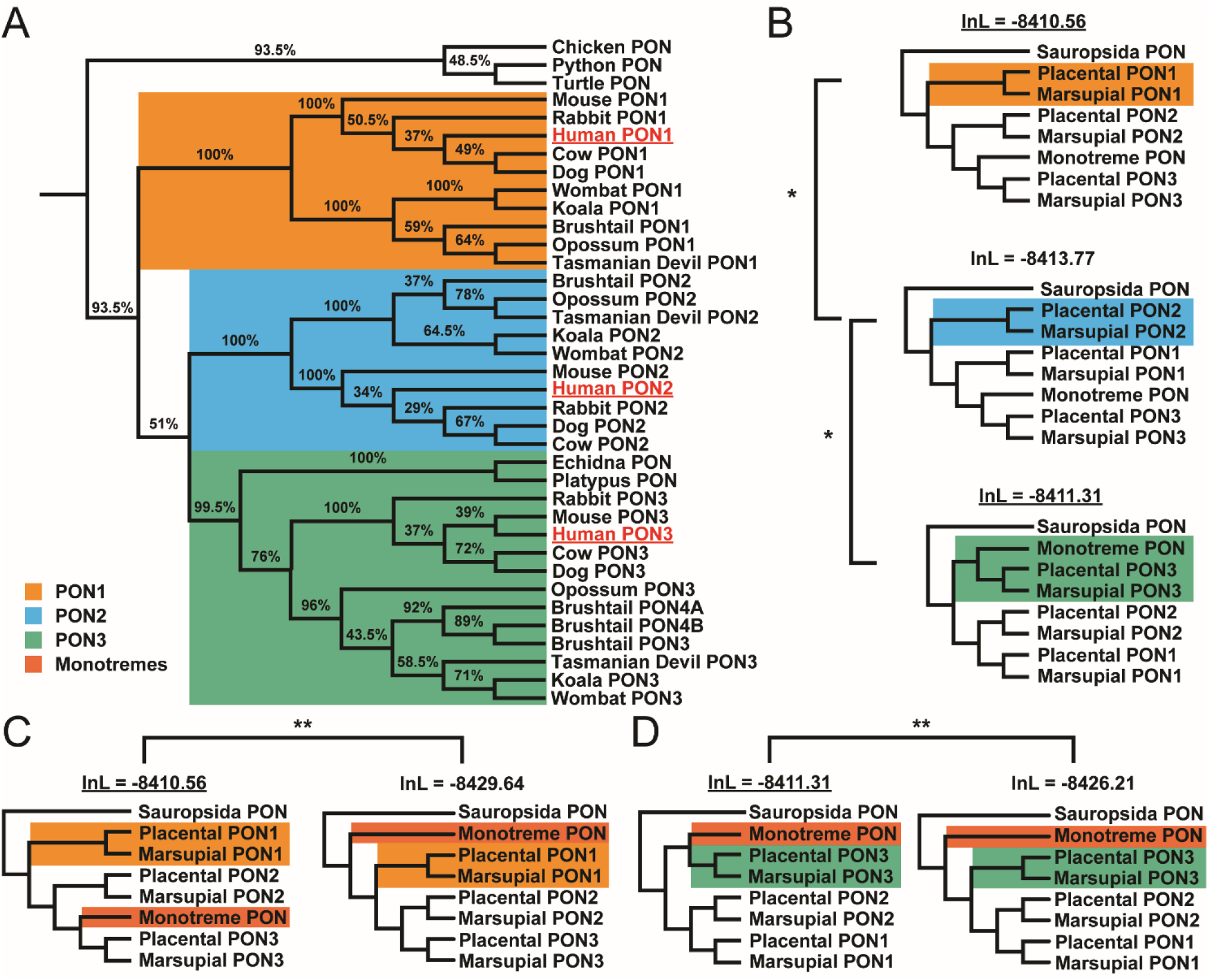
Evolution of PON in tetrapods. (A) Phylogenetic tree of PON family proteins in tetrapods determined by PhyML based on multiple sequence alignment. Bootstrap support values are shown as percentages out of 200. Brushtail PON4 was split into two separate genes based on RNA-seq evidence. Human sequences were underlined to help orient the reader. (B) Phylogenetic tree models represent the scenario in which each of the PONs is ancestral compared to the other two. * indicates p-value was less than 0.05. The underlined log likelihood indicates the significantly better model. (C,D) Two sets of models compare the placement of the monotremes either basally along with birds and lizards (i.e. sauropsida) or within PON3 with either PON1 being ancestral (C) or PON3 being ancestral (D). ** indicates p-value < 1e-8. The underlined log likelihood indicates the significantly better model. Rooted and unrooted trees produced the same log likelihood score.

In addition to instances of *PON* expansion, there is also evidence of *PON* loss. Upon examining multiple species in the same taxon without a *PON* gene, this leads to the possibility of ancestral losses of *PON* in the ctenophores, placozoans, urochordates, arthropods, and platyhelminths (Fig 4, Fig S3). Outside those lineages, there are several parasitic species (i.e. *Soboliphyme baturini, Teladorsagia circumcincta, Hirudo medicinalis*, Myxozoan cnidarians), in which we were unable to identify a single *PON* gene. This was not surprising as species which evolve to become parasitic or symbiotic tend to undergo genome reduction(Wolf and Koonin 2013).

### PON expansion occurred after divergence of mammals from sauropsida

RefSeq-identified PON protein sequences from fifteen tetrapod species were collected(Pruitt et al. 2005). Each of the five placental and five marsupial mammals encoded three PON proteins. Both monotreme mammals as well as the three sauropsid (reptile and birds) species each have a single copy of *PON*. While simple parsimony of gene counts would suggest that the duplications leading to three therian *PON*s occurred after divergence from monotremes, our phylogenetic analysis reveals a strong clustering of three distinct mammalian *PON* clusters (Fig 2A). Importantly, the monotreme *PON* genes clustered in the marsupial and placental *PON3* sequences with high bootstrap support (99.5%), instead of falling outside the PON gene duplications. This indicates that the mammalian *PON* gene family expanded to at least three members after the divergence of mammals from sauropsida (the ancestor to reptiles and birds) but before divergence of monotremes. This is further bolstered by the evidence of distinct clusters corresponding to the divergence of monotremes, marsupials, and placentals within each of the individual *PON* groups with 100% bootstrap support except for the marsupial *PON3* cluster which has 96% bootstrap support. The lack of additional monotreme *PON* genes strongly suggests that the ancestor of extant monotremes lost its *PON1* and *PON2* genes or the ancestor to these two genes after it diverged from the therian ancestor.

To determine which of the three *PON*s diverged first, we considered three separate models in which each of the mammalian *PON*s diverged before the other two (Fig 2B). Using the multiple protein alignment from the first analysis, PAML determined the likelihood that the alignment would support that model. The model with the highest likelihood shows *PON1* diverging first (-8410.56), although it was not significantly better than the model with *PON3* diverging first (-8411.31). Thus, we cannot be certain which of those two *PON*s is ancestral to the other (p-value = 0.22); however, it can be stated with statistical significance that *PON2* did not diverge before *PON1* (-8413.77, -8410.56, p-value = 0.0113) nor did *PON2* diverge before *PON3* (-8413.77, -8411.31, p-value = 0.0267) (Fig 2B).

To confirm that the monotreme *PONs* belong with the ancestral *PON3* group, two additional models were created. Both models clustered the monotremes *PON*s with the sauropsida (birds and reptiles) *PON*s instead of mammalian *PON3*, but one of them had *PON1* diverged first (Fig 2C) while the other had *PON3* diverge first (Fig 2D). Regardless of whether *PON1* or *PON3* diverged before the other, the models in which the monotremes’ *PON*s belong to the mammalian *PON3* group were strongly preferred (P-values = 6.5×10^−10^ and 4.81×10^−8^). This indicates they lost either *PON1* and *PON2* separately or the ancestor to both *PON1* and *PON2*.

### Case Study: Recent *PON* expansion under positive selection in brushtail possum

In addition to the three *PON*s that were expected to be found in the brushtail possum, a marsupial mammal, BLAST identified another locus (XP_036615431.1) in between the brushtail *PON3* and *PON1*. Upon closer examination this single locus in brushtail contained what could be two new *PON* genes. We therefore divided this locus into two halves, each forming a complete arylesterase domain with typical PON gene structure, to form *PON4A* and *PON4B*, since RNA-seq reads show they are transcribed independently (Fig 3A). PON4A is comprised of annotated residues 1-337 of XP_036615431.1, and PON4B consists of residues 338-717, although this study suggests the gene model should be updated as two separate genes. Based on sequence similarity, both *PON4A* and *PON4B* seemed to be expansions from the brushtail PON3 gene (Fig 2A).

**Figure 3.**
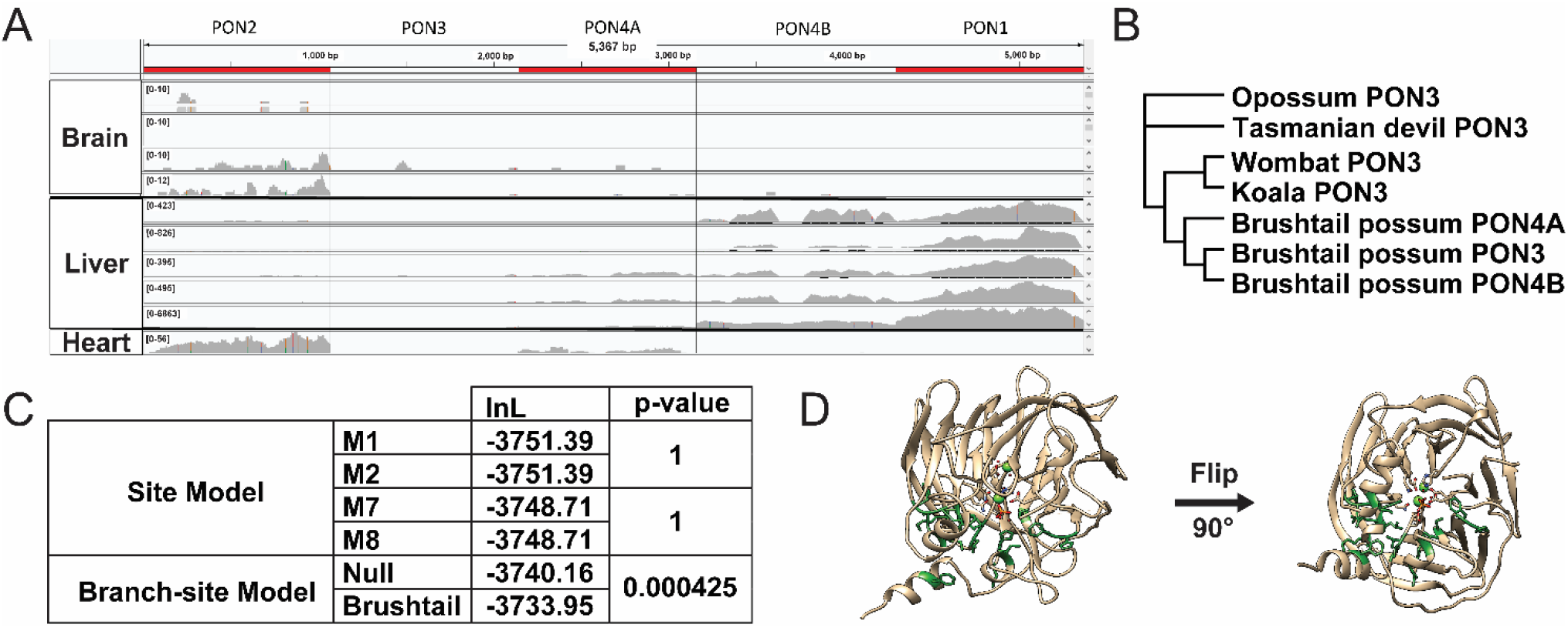
Brushtail PON3 and recent PON4A/B under positive selection. (A) Screenshot from IGV showing a mapped reads to a concatenated transcriptome for all brushtail possum *PON* genes. Tissue samples include four brain, five liver, and one heart sample. The order of the concatenated PON genes is listed above the screenshot. Order was chosen to mimic the chromosomal order found in the brushtail genome. (B) Phylogenetic tree of marsupial *PON3* genes used for PAML and BUSTED analysis shown. (C) Log likelihood values were determined by PAML. The marsupial tree in figure 3B was used in the marsupial analysis (D) 3D protein image of rabbit PON1 (PDB:1V04). Corresponding sites under positive selection in brushtail are colored green and show their atomic structure. Residues which are part of the active site also show their atomic structure.

To verify that *PON4A* and *PON4B* were real and not the result of an assembly error, brushtail possum RNA-seq reads were mapped to the five *PON* mRNA sequences (Fig 3A). Ten RNA-seq samples were available with four from the brushtail brain, five from the liver, and one from the heart. In the brain, there was a low level of *PON2* expression detected. As anticipated, there was robust expression of *PON1* in the liver as observed in other therian mammals. There was no *PON3* expression in the liver as was expected given PON3’s liver expression in other therian mammals. Instead, there seemed to be expression of an isoform of *PON4A* in the liver. In the heart, there was noted expression of *PON2* as was expected, and surprisingly, there was also low expression of *PON4B*.

Given the unexpected lack of expression of *PON3* and tissue-specific expression of PON4A and PON4B, we next looked to see if there was evidence of positive selection associated with this recent expansion and diversification of brushtail *PON3* into *PON4A* and *PON4B*. We first used CODEML from the PAML package to compare sites models, M1 vs M2 and M7 vs M8, to test if there was positive selection within the entire marsupial PON3 clade (Fig 3B, 3C), and we found no evidence for it. We then tested if there was evidence of episodic positive selection associated with the brushtail lineage and its gene duplications compared to the rest of the marsupial *PON3* genes using branch-site models. Indeed, we observed that the branch-site model allowing positive selection in the brushtail *PON*s fit the data significantly better than the null model (p-value = 0.000425), and it was estimated that 11.5% of the positions in brushtail *PON3* sequences evolved under positive selection. The subsequent Bayes

Empirical Bayes (BEB) analysis revealed 19 of the 352 positions under positive selection with a posterior probability exceeding 0.5. Sixteen of the sites were mapped to the rabbit PON1 structure (PDB 1V04) so we could visualize where within the *PON* protein the positive selection was occurring. We observed the residues to be clustered around the catalytic active site (Fig 3D) and we determined that these sites under positive selection are clustered together more closely than would be expected by chance (permutation p-value < 1e-6). These changes near the active site could have increased the specificity of these enzymes for a specific yet unidentified substrate(s) which enhanced the fitness of this species.

As an additional method to test for positive selection occurring within the brushtail species, we ran Branch-site Unrestricted Statistical Test for Episodic Diversification (BUSTED, http://www.datamonkey.org/busted). While it found that the unconstrained model (logL = -3726.8) with the brushtail *PON3, PON4A*, and *PON4B* as the foreground sequences fit the data better than the null model (logL = -3728.1), it did not reject the null at an alpha of 0.05 (p-value 0.142), so inferences of positive selection should be treated with some caution. A major difference between these models and those of CODEML is that BUSTED accommodates variation in rates of synonymous site divergence.

## Discussion

Through this phylogenetic analysis of the *PON* gene family, we see the changes in *PON* copy number are not restricted to just mammals and cannot be explained as the result of the whole genome duplication in vertebrates and teleosts, allowing for future investigation into common selective pressures which favor the expansion or reduction in the number of PON members. While most mammals have three *PON* genes, the process of *PON1* becoming a pseudogene within diving mammals has raised the question of when this gene family expanded during evolutionary time. With the genomes of two monotremes, we concluded that the mammalian *PON* expansion occurred before the divergence of monotremata from theria, in the ancestral lineage leading to all mammals. This prompted a more extensive investigation of *PON* genes throughout all metazoa which revealed that there have been multiple independent expansions and contractions of *PON* throughout metazoa. Finally, a closer investigation of the brushtail possum genome revealed there has been a local expansion of *PON3* within that species associated with positively selected amino acid changes and rapid divergence in expression patterns across tissues.

In contrast to previous findings, the results presented in this paper suggest that PON1 or PON3 diverged before *PON2* (Fig 2). The initial study was perhaps limited by the use of four placental mammals(Draganov and La Du 2004). In a more recent study, which used six placental mammals, *PON3* was identified as likely being ancestral to *PON1* and *PON2*(Bar-Rogovsky et al. 2013). In this study comprised of five placental mammals, five marsupials, and two monotremes, we are unable to conclude if *PON1* or *PON3* diverged first; however, we can conclude that *PON2* did not diverge first. Because the monotreme sequences cluster better with *PON3* instead of diverging from the ancestral branch leading to all mammals, this informs us that the *PON* family expanded before monotremes diverged. Given the lack of *PON1* and *PON2* in the otherwise contiguous monotreme assemblies this strongly suggests that monotremes had *PON1* and *PON2* or their ancestor but then lost them.

Throughout the metazoan tree, there are multiple examples of *PON* expansion and several independent suggestions of PON loss (Fig 4, Fig S3). Given the promiscuous nature of these enzymes, it can be difficult to determine what evolutionary pressures favored expansion and reduction of the *PON* gene family. There could be different functions of these enzymes which are favored in each independent instance. One possible selection pressure is a change in response to oxidative stress since it has been proposed that *PON1* at least has a role in mitigating oxidative damage to lipids(Aviram et al. 2000). Oxidative stress management is different in aquatic environments compared to living on land because diving mammals need to tolerate repeated diving-induced ischemia and reperfusion(Allen and Vázquez-Medina 2019). Given the convergent loss of a functional *PON1* gene in aquatic mammals, this suggests that its loss is beneficial for increasing marine mammals tolerance of repeated ischemia and reperfusion(Meyer et al. 2018). Although it is not clear why losing an enzyme that is thought to mitigate the effects of oxidative stress would be lost in species that encounter increased oxidative stress.

**Figure 4.**
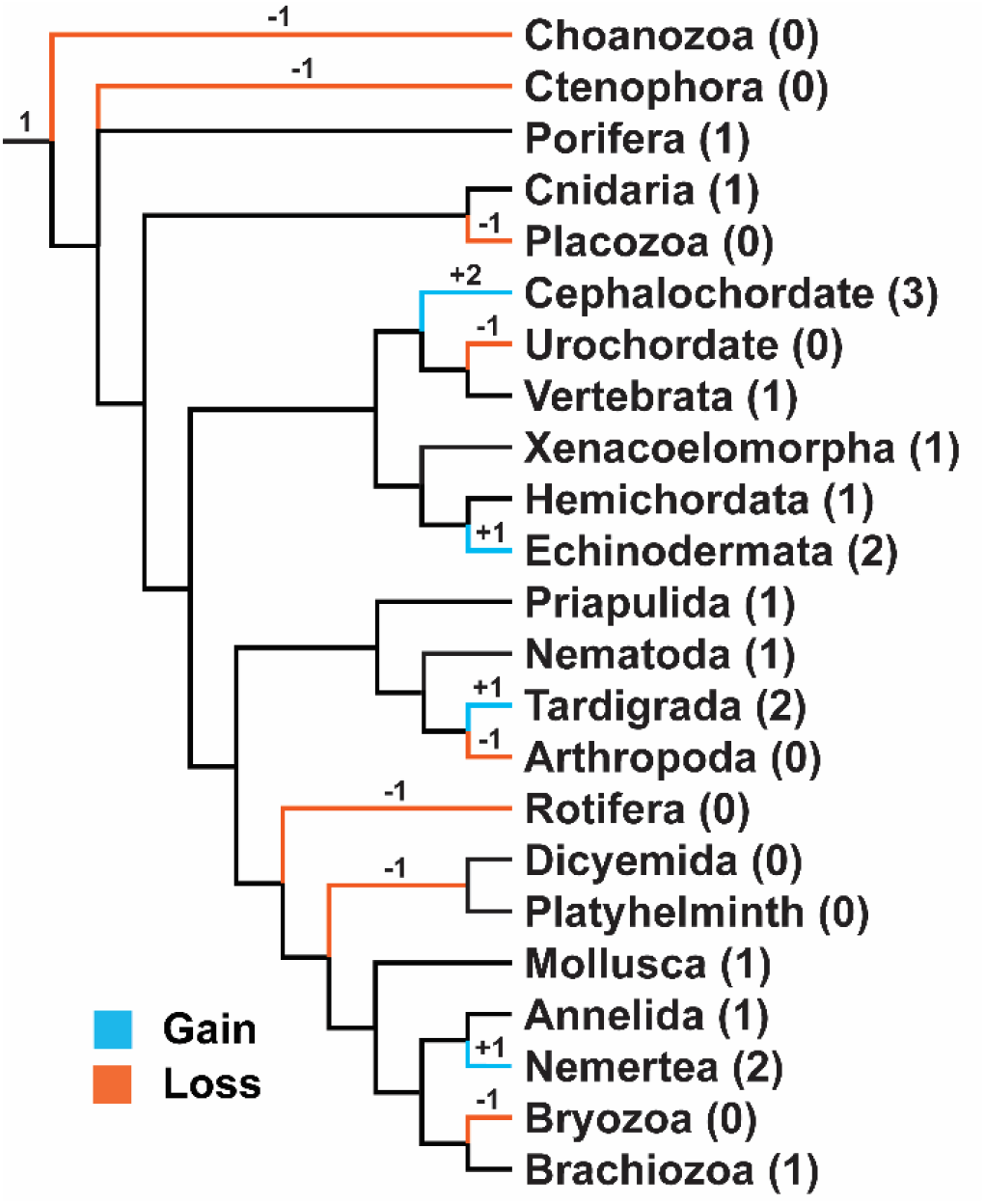
Overview of observed changes in the number of PON genes throughout metazoa. Phylogenetic tree represent the approximate time when changes in the number of PON genes occurred in metazoan broadly(Laumer et al. 2019; Philippe et al. 2019; Kapli and Telford 2020). More detailed phylogenetic tree is available as supplemental figure 3. The relative timing of the changes is indicated by the number and sign above the branch. Number in between parenthesis after the phylum name indicates the number of ancestral genes which could be detected for the terminal branch. The lack of changes noted in the deeper portions of this tree are not meant to indicate that there was no change in gene copy number in those branches. Rather, it is merely a reflection of the limitation of this study to probe those branches. Ctenophora and Porifera are shown as a polytomy as their exact relationship has not yet been elucidated(Laumer et al. 2019; Li et al. 2021; Redmond and McLysaght 2021). Placement of Dicyemida is not well resolved at the time of this publication(Zverkov et al. 2019).

Resistance to bacteria is another possible selection pressure behind the dynamic evolution of the *PON* family. One way bacteria progress as an infection is through the construction of a biofilm(Smith and Iglewski 2003). A common signaling molecule bacteria use is homoserine lactone (HSL) which PON2 has shown evidence of being able to degrade(Smith and Iglewski 2003; Bar-Rogovsky et al. 2013). This could potentially explain the large *PON* expansions observed within cephalochordates, ambulacraria, and bivalves. Members within these taxonomical groups feed primarily by filtering nutrients from water.

Inhibiting biofilm formation would be important for these species so that the biofilm does not inhibit their ability to extract nutrients and sustenance from the water. Another possible explanation for the large *PON* expansion observed in these water filtering species is they are the frequent recipients of multiple horizontal transfer from bacteria given their close contact with bacteria.

While two selective pressures have been offered as explanation for *PON* expansions, neither yet sufficiently explains the brushtail possum specific duplication of *PON3*. While an increased bacterial burden in the brushtail possum could be an explanation, an expansion of *PON2* would be more likely as PON2 is more efficient at hydrolyzing HSLs compared to PON3, at least in other theria. This suggests there are more selective pressures related to *PON3* which promoted the fixation and divergence of these duplications in the brushtail possum genome. Further sequencing of other brushtail possum subspecies and related species could determine when this expansion took place and provide clues as to what selective pressures favored it. Additionally, more RNA-seq samples from different tissue types are needed to determine where each of the *PON* genes are being expressed. Brushtail *PON4B* and *PON1* have been shown to be expressed in the liver while *PON2* and *PON4A* are observed in the heart. It is not clear in what tissue (if any) brushtail *PON3* is expressed.

From the results of these studies, it is fair to propose that this promiscuous family of enzymes plays an ever-changing role depending on lineage. While we can theorize why this gene family expanded in some lineages and contracted in others, additional experiments are needed to test these theories. Certainly, the expansion of *PON3* within the brushtail possum hints that there are still other explanations waiting to be discovered.

## Data Availability

The protein models underlying this article are individually available at a variety of repositories Zenodo(Copley et al. 2018; Kvist et al. 2019; schultz and Francis 2020), National Center for Biotechnology Information (NCBI), Ensembl, SIMRBASE(Data), OIST Marine Genome Projects(Data), Github(Ryan), Harvard Dataverse(Qingxiang), Plos One(Delroisse et al. 2016), Google Drive(Data), Neurobase(Data), Bitbucket, Dryad(Data), Ephybase(Data), Reefgenomics(Liew et al. 2016), GigaDB(Data), National Genomics Data Center (NGDC)(Chen et al. 2021), Figshare(Data), PeerJ(Jin et al. 2020), Planmine(Rozanski et al. 2019), NHGRI(Data), and the Ryan Lab website(Ryan). All protein models have their assembly information listed in Supplementary Table 1. In addition, they are designated as having come from NCBI, Ensembl or with URL listed. All newick trees and multiple sequence alignments underlying this article are available at the Clark website https://clark.genetics.utah.edu/software-data-and-collaborators/. The protein crystal structure is available at https://www.rcsb.org/structure/1V04.

## Acknowledgement

The support and resources from the Center for High Performance Computing at the University of Utah are gratefully acknowledged. The computational resources used were partially funded by the NIH Shared Instrumentation Grant 1S10OD021644-01A1. This work was supported by the National Institutes of Health (grant numbers 3T32DK007115 to AMG, K99GM144774 to AMG, R01 HG009299 to AMG and NLC, and R01 EY030546 to SAML, JSP, and NLC). We thank the rest of the Clark lab, Paige Eberle, and Chris Stringham for thoughtful suggestions and useful discussions.

