## Supplemental Table 2 for "Highly Dynamic Gene Family Evolution Suggests Changing Roles for *PON* Genes Within Metazoa"

| | | | | | | PON1 | | | PON2 | | | PON3 | |
|---|---|---|---|---|---|---|---|---|---|---|---|---|---|
| Common Name | Species | Group | PON name | RefSeq accession | Query Cover | E value | Per. Ident | Query Cover | E value | Per. Ident | Query Cover | E value | Per. Ident |
| Chicken | Gallus gallus | Outgroup | pon | NP_001188397.1 | 93% | 9.E-174 | 69.58% | 93% | 0.E+00 | 74.62% | 93% | 4.E-171 | 66.77% |
| Painted Turtle | Chrysemys picta | Outgroup | pon | XP_005299911.1 | 100% | 0.E+00 | 67.32% | 100% | 0.E+00 | 74.01% | 100% | 2.E-180 | 67.51% |
| Python | Python bivittatus | Outgroup | pon | XP_007425171.1 | 100% | 2.E-162 | 61.52% | 100% | 0.E+00 | 67.61% | 100% | 1.E-164 | 61.69% |
| Short-beaked Echidna | Tachyglossus aculeatus | Monotremata | pon | XP_038598090.1 | 100% | 2.E-142 | 56.06% | 94% | 1.E-148 | 60.12% | 100% | 6.E-153 | 57.91% |
| Platypus | Ornithorhynchus anatinus | Monotremata | pon | XP_028927165.1 | 93% | 3.E-143 | 56.46% | 94% | 4.E-153 | 60.42% | 94% | 7.E-159 | 60.12% |
| Gray short-tailed opossum | Monodelphis domestica | Marsupilia | pon1 | XP_007505543.1 | 93% | 5.E-176 | 69.79% | 93% | 4.E-162 | 65.26% | 93% | 2.E-141 | 58.01% |
| Gray short-tailed opossum | Monodelphis domestica | Marsupilia | pon2 | XP_007506048.1 | 93% | 2.E-164 | 65.77% | 94% | 0.E+00 | 77.08% | 94% | 8.E-159 | 64.29% |
| Gray short-tailed opossum | Monodelphis domestica | Marsupilia | pon3 | XP_007506039.1 | 100% | 6.E-150 | 57.70% | 100% | 6.E-160 | 60.67% | 100% | 1.E-171 | 65.73% |
| Tasmanian Devil | Sarcophilus harrisii | Marsupilia | pon1 | XP_031796628.1 | 92% | 0.E+00 | 74.01% | 91% | 4.E-162 | 66.26% | 92% | 9.E-145 | 59.94% |
| Tasmanian Devil | Sarcophilus harrisii | Marsupilia | pon2 | XP_031796625.1 | 93% | 3.E-168 | 66.16% | 93% | 0.E+00 | 78.48% | 93% | 2.E-159 | 63.94% |
| Tasmanian Devil | Sarcophilus harrisii | Marsupilia | pon3 | XP_031796627.1 | 93% | 1.E-142 | 58.26% | 95% | 1.E-161 | 62.83% | 95% | 3.E-171 | 66.96% |
| Common Wombat | Vombatus ursinus | Marsupilia | pon1 | XP_027704510.1 | 94% | 0.E+00 | 71.94% | 94% | 2.E-166 | 67.36% | 94% | 2.E-143 | 58.75% |
| Common Wombat | Vombatus ursinus | Marsupilia | pon2 | XP_027704673.1 | 100% | 7.E-174 | 65.07% | 100% | 0.E+00 | 81.92% | 100% | 2.E-166 | 63.28% |
| Common Wombat | Vombatus ursinus | Marsupilia | pon3 | XP_027704640.1 | 93% | 9.E-149 | 60.06% | 95% | 5.E-163 | 64.01% | 95% | 5.E-176 | 69.44% |
| Common Brushtail possum | Trichosurus vulpecula | Marsupilia | pon1 | XP_036615148.1 | 94% | 0.E+00 | 74.63% | 94% | 2.E-176 | 70.03% | 94% | 5.E-150 | 60.24% |
| Common Brushtail possum | Trichosurus vulpecula | Marsupilia | pon2 | XP_036617064.1 | 100% | 7.E-168 | 64.23% | 100% | 0.E+00 | 80.79% | 100% | 5.E-164 | 63.28% |
| Common Brushtail possum | Trichosurus vulpecula | Marsupilia | pon3 | XP_036615435.1 | 100% | 1.E-144 | 56.23% | 100% | 1.E-160 | 60.56% | 100% | 3.E-165 | 63.33% |
| Common Brushtail possum | Trichosurus vulpecula | Marsupilia | pon4A | XP_036615431.1 | 100% | 3.E-152 | 60.67% | 100% | 1.E-164 | 63.38% | 100% | 7.E-175 | 67.89% |
| Common Brushtail possum | Trichosurus vulpecula | Marsupilia | pon4B | XP_036615431.1 | 100% | 3.E-152 | 60.67% | 100% | 1.E-164 | 63.38% | 100% | 7.E-175 | 67.89% |
| Koala | Phascolarctos cinereus | Marsupilia | pon1 | XP_020855645.1 | 100% | 0.E+00 | 72.19% | 100% | 1.E-169 | 65.73% | 99% | 5.E-144 | 57.10% |
| Koala | Phascolarctos cinereus | Marsupilia | pon2 | XP_020855737.1 | 100% | 2.E-171 | 64.79% | 100% | 0.E+00 | 82.49% | 100% | 4.E-165 | 62.99% |
| Koala | Phascolarctos cinereus | Marsupilia | pon3 | XP_020855646.1 | 100% | 1.E-151 | 57.46% | 100% | 5.E-168 | 62.15% | 100% | 0.E+00 | 68.64% |
| Mouse | Mus musculus | Placentalia | pon1 | NP_035264.2 | 100% | 0.E+00 | 82.25% | 100% | 9.E-175 | 64.51% | 100% | 4.E-157 | 59.72% |
| Mouse | Mus musculus | Placentalia | pon2 | NP_899131.1 | 93% | 6.E-168 | 66.37% | 100% | 0.E+00 | 88.14% | 100% | 2.E-165 | 64.41% |
| Mouse | Mus musculus | Placentalia | pon3 | NP_766594.1 | 100% | 6.E-154 | 57.18% | 100% | 2.E-170 | 62.99% | 100% | 0.E+00 | 81.36% |
| Rabbit | Oryctolagus cuniculus | Placentalia | pon1 | NP_001075766.1 | 100% | 0.E+00 | 85.63% | 93% | 8.E-170 | 66.67% | 93% | 9.E-152 | 60.36% |
| Rabbit | Oryctolagus cuniculus | Placentalia | pon2 | XP_002713838.1 | 100% | 9.E-179 | 65.35% | 100% | 0.E+00 | 93.22% | 100% | 3.E-175 | 64.97% |
| Rabbit | Oryctolagus cuniculus | Placentalia | pon3 | NP_001075547.1 | 93% | 7.E-150 | 59.76% | 95% | 3.E-167 | 65.49% | 95% | 0.E+00 | 84.37% |
| Cow | Bos taurus | Placentalia | pon1 | NP_001039734.1 | 100% | 0.E+00 | 82.25% | 92% | 2.E-167 | 65.96% | 93% | 5.E-150 | 59.16% |
| Cow | Bos taurus | Placentalia | pon2 | NP_001013606.1 | 93% | 3.E-171 | 67.87% | 100% | 0.E+00 | 93.22% | 100% | 1.E-169 | 66.10% |
| Cow | Bos taurus | Placentalia | pon3 | NP_001068947.1 | 93% | 5.E-150 | 58.86% | 100% | 1.E-169 | 66.38% | 100% | 0.E+00 | 81.36% |
| Dog | Canis lupus familiaris | Placentalia | pon1 | XP_850219.2 | 100% | 0.E+00 | 87.61% | 92% | 9.E-166 | 66.57% | 93% | 5.E-149 | 59.46% |
| Dog | Canis lupus familiaris | Placentalia | pon2 | NP_001003205.2 | 93% | 4.E-169 | 67.27% | 100% | 0.E+00 | 93.22% | 94% | 2.E-164 | 65.18% |
| Dog | Canis lupus familiaris | Placentalia | pon3 | XP_038291432.1 | 93% | 2.E-154 | 62.24% | 93% | 1.E-162 | 66.06% | 93% | 0.E+00 | 82.12% |
| Human | Homo sapiens | Placentalia | pon1 | NP_000437.3 | 100% | 0.E+00 | 100.00% | 100% | 9.E-178 | 65.92% | 100% | 2.E-161 | 60.85% |
| Human | Homo sapiens | Placentalia | pon2 | NP_000296.2 | 100% | 4.E-172 | 65.92% | 100% | 0.E+00 | 100.00% | 100% | 5.E-169 | 65.54% |
| Human | Homo sapiens | Placentalia | pon3 | NP_000931.1 | 100% | 1.E-154 | 60.85% | 100% | 1.E-168 | 65.54% | 100% | 0.E+00 | 100.00% |
