## Supplemental Table 3 for "Highly Dynamic Gene Family Evolution Suggests Changing Roles for *PON* Genes Within Metazoa"

| Sample | NCBI_Accession | Tissue # | Layout | Read-length (bp) | Filtered Reads | Genome Reads | Transcript |
|---|---|---|---|---|---|---|---|
| 1 | ERR4083814 | Metagenome 1 | Paired | 150 | 2191193 | 26 | 0 |
| 2 | ERR4083805 | Metagenome 2 | Paired | 150 | 2475430 | 204 | 2 |
| 3 | ERR4083613 | Metagenome 3 | Paired | 150 | 1557425 | 358 | 0 |
| 4 | ERR4083612 | Metagenome 4 | Paired | 150 | 1504977 | 66 | 1 |
| 5 | SRR11483671 | Liver 1 | Single | 50 | 41960633 | 690100 | 9 |
| 6 | SRR11483672 | Liver 2 | Single | 50 | 48578493 | 993502 | 0 |
| 7 | SRR11483673 | Liver 3 | Single | 50 | 42457267 | 466352 | 41 |
| 8 | SRR11483674 | Liver 4 | Single | 50 | 48980015 | 264593 | 64 |
| 9 | SRR11481814 | Brain 1 | Single | 50 | 42068707 | 309990 | 7366 |
| 10 | SRR11481815 | Brain 2 | Single | 50 | 47159144 | 219920 | 9634 |
| 11 | SRR11481816 | Brain 3 | Single | 50 | 40945204 | 220466 | 6263 |
| 12 | SRR11481817 | Brain 4 | Single | 50 | 44994063 | 229284 | 8160 |
| 13 | SRR8658969 | Liver 5 | Paired | 101 | 47538674 | 214045 | 84169 |
| 14 | SRR8658968 | Heart | Paired | 101 | 58087750 | 104862 | 388 |
