## Supplemental Figures for "Highly Dynamic Gene Family Evolution Suggests Changing Roles for *PON* Genes Within Metazoa"

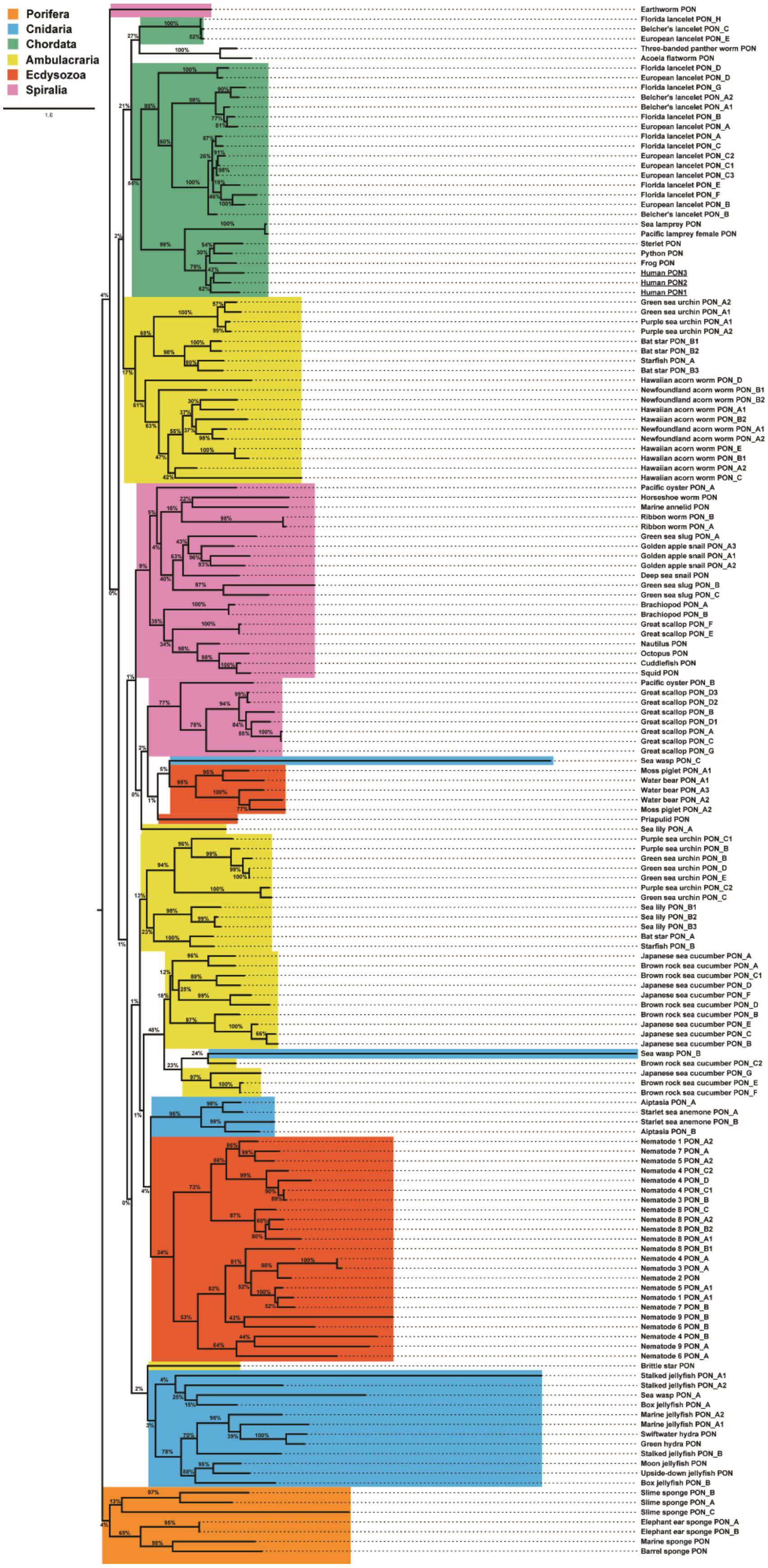
Porifera
Cnidaria
Chordata
Ambulacraria
Ecdysozoa
Spiralia
1.0
Earthworm PON
Florida lancelet PON_H
Belcher's lancelet PON_C
European lancelet PON_E
Three-banded panther worm PON
Acoela flatworm PON
Florida lancelet PON_D
European lancelet PON_D
Florida lancelet PON_G
Belcher's lancelet PON_A2
Belcher's lancelet PON_A1
Florida lancelet PON_B
European lancelet PON_A
Florida lancelet PON_A
Florida lancelet PON_C
European lancelet PON_C2
European lancelet PON_C1
European lancelet PON_C3
Florida lancelet PON_E
Florida lancelet PON_F
European lancelet PON_B
Belcher's lancelet PON_B
Sea lamprey PON
Pacific lamprey female PON
Sterlet PON
Python PON
Frog PON
Human PON3
Human PON2
Human PON1
Green sea urchin PON_A2
Green sea urchin PON_A1
Purple sea urchin PON_A1
Purple sea urchin PON_A2
Bat star PON_B1
Bat star PON_B2
Starfish PON_A
Bat star PON_B3
Hawaiian acorn worm PON_D
Newfoundland acorn worm PON_B1
Newfoundland acorn worm PON_B2
Hawaiian acorn worm PON_A1
Hawaiian acorn worm PON_B2
Newfoundland acorn worm PON_A1
Newfoundland acorn worm PON_A2
Hawaiian acorn worm PON_E
Hawaiian acorn worm PON_B1
Hawaiian acorn worm PON_A2
Hawaiian acorn worm PON_C
Pacific oyster PON_A
Horseshoe worm PON
Marine annelid PON
Ribbon worm PON_B
Ribbon worm PON_A
Green sea slug PON_A
Golden apple snail PON_A3
Golden apple snail PON_A1
Golden apple snail PON_A2
Deep sea snail PON
Green sea slug PON_B
Green sea slug PON_C
Brachiopod PON_A
Brachiopod PON_B
Great scallop PON_F
Great scallop PON_E
Nautilus PON
Octopus PON
Cuddlefish PON
Squid PON
Pacific oyster PON_B
Great scallop PON_D3
Great scallop PON_D2
Great scallop PON_B
Great scallop PON_D1
Great scallop PON_A
Great scallop PON_C
Great scallop PON_G
Sea wasp PON_C
Moss piglet PON_A1
Water bear PON_A1
Water bear PON_A3
Water bear PON_A2
Moss piglet PON_A2
Priapulid PON
Sea lily PON_A
Purple sea urchin PON_C1
Purple sea urchin PON_B
Green sea urchin PON_B
Green sea urchin PON_D
Green sea urchin PON_E
Purple sea urchin PON_C2
Green sea urchin PON_C
Sea lily PON_B1
Sea lily PON_B2
Sea lily PON_B3
Bat star PON_A
Starfish PON_B
Japanese sea cucumber PON_A
Brown rock sea cucumber PON_A
Brown rock sea cucumber PON_C1
Japanese sea cucumber PON_D
Japanese sea cucumber PON_F
Brown rock sea cucumber PON_D
Brown rock sea cucumber PON_B
Japanese sea cucumber PON_E
Japanese sea cucumber PON_C
Japanese sea cucumber PON_B
Sea wasp PON_B
Brown rock sea cucumber PON_C2
Japanese sea cucumber PON_G
Brown rock sea cucumber PON_E
Brown rock sea cucumber PON_F
Aiptasia PON_A
Starlet sea anemone PON_A
Starlet sea anemone PON_B
Aiptasia PON_B
Nematode 1 PON_A2
Nematode 7 PON_A
Nematode 5 PON_A2
Nematode 4 PON_C2
Nematode 4 PON_D
Nematode 4 PON_C1
Nematode 3 PON_B
Nematode 8 PON_C
Nematode 8 PON_A2
Nematode 8 PON_B2
Nematode 8 PON_A1
Nematode 8 PON_B1
Nematode 4 PON_A
Nematode 3 PON_A
Nematode 2 PON
Nematode 5 PON_A1
Nematode 1 PON_A1
Nematode 7 PON_B
Nematode 9 PON_B
Nematode 6 PON_B
Nematode 4 PON_B
Nematode 9 PON_A
Nematode 6 PON_A
Brittle star PON
Stalked jellyfish PON_A1
Stalked jellyfish PON_A2
Sea wasp PON_A
Box jellyfish PON_A
Marine jellyfish PON_A2
Marine jellyfish PON_A1
Swiftwater hydra PON
Green hydra PON
Stalked jellyfish PON_B
Moon jellyfish PON
Upside-down jellyfish PON
Box jellyfish PON_B
Slime sponge PON_B
Slime sponge PON_A
Slime sponge PON_C
Elephant ear sponge PON_A
Elephant ear sponge PON_B
Marine sponge PON
Barrel sponge PON

Supplementary Figure 1 Evolution of PON in metazoans with branch lengths. Phylogenetic tree of PON family proteins within metazoa as determined by RAxML based on multiple sequence alignment. The numbers on the branches indicate the branch length as well as the relative lengths of the actual branch lengths. If species have multiple PON genes and they are located on different chromosome/scaffold, then they are indicated by different alphabet characters. If they PON genes are located on the same chromosome/scaffold and are within one megabase of another PON gene, this is indicated by a number.

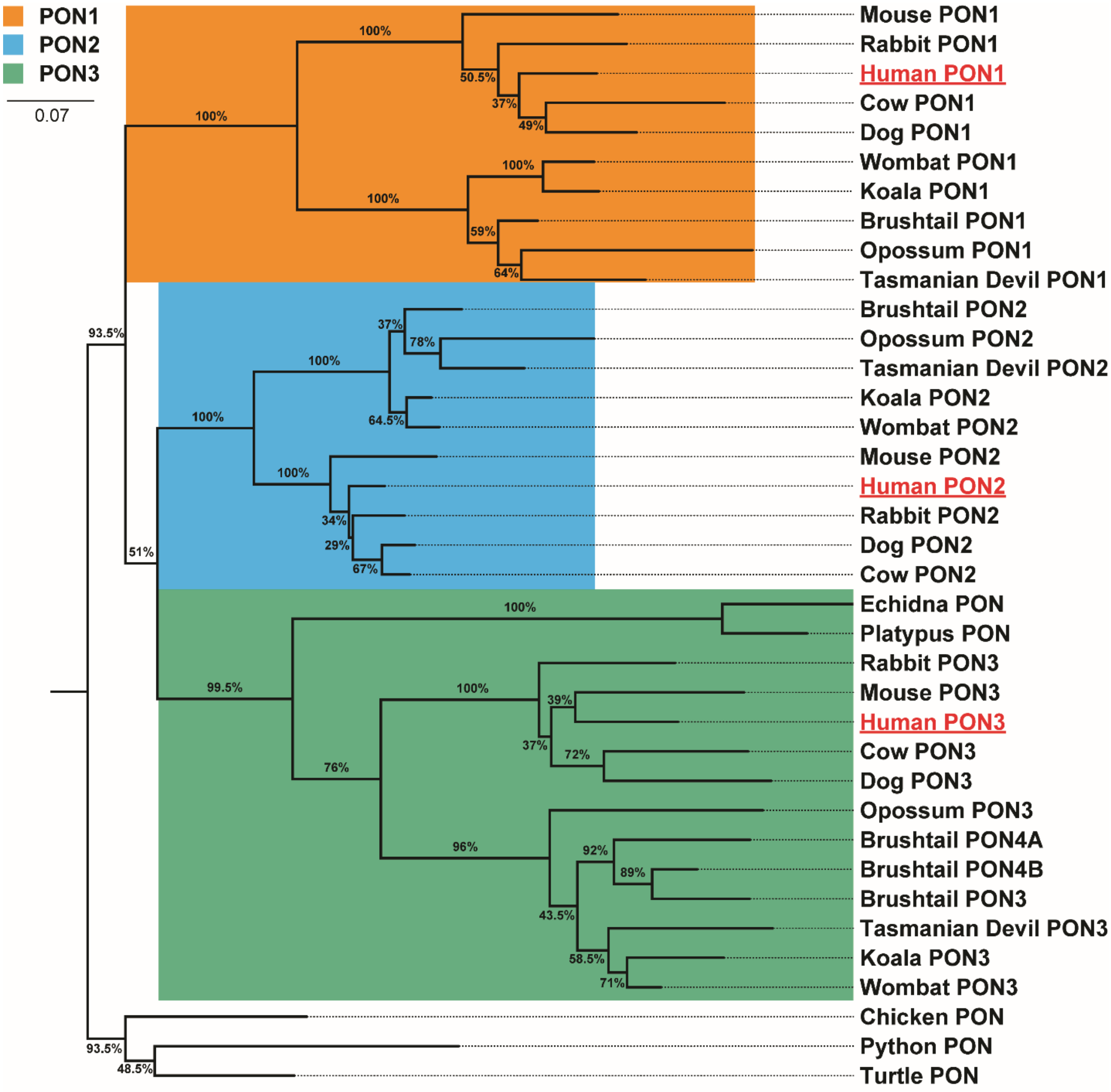


Supplementary Figure 2 Evolution of PON in tetrapods with branch lengths. This phylogenetic tree has the same topology, or overall shape as the phylogenetic tree in figure 2A, and the values above the branches indicate the bootstrap values as a percent out of 200. The branch lengths have been scaled to reflect the relative branch lengths between all the sequences. Human sequences were underlined for ease of orienting the reader.

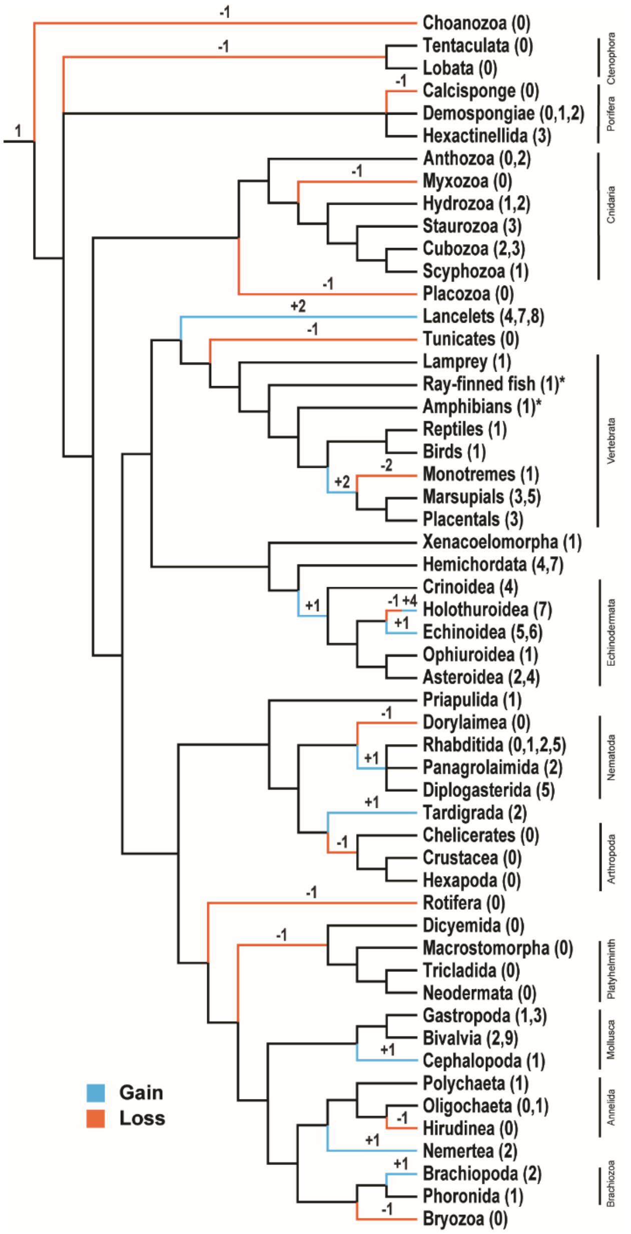


Supplementary Figure 3 Observed changes in number of *PON* genes in different clades. Cladogram represents relationships between these taxonomical groups[58,63–70]. Phyla for which there is more sequences available (i.e. vertebrates) are broken down into more granular taxa. If the phylum name was not used, then it is indicated on the right-hand side. Each clade represents one to five species studied. The relative timing of the changes is indicated by the number and sign above the branch. Number in between parenthesis after the clade name indicates the number of *PON* genes detected for the terminal branch. Numbers which do not add up with the changes indicate local changes observed within a subset of species in that taxon. Taxon with an asterisk (*) have members which are known to have undergone a recent whole genome duplication. Those members were not investigated in this study. The relationships within Porifera[71] and Nematoda[72] are currently unresolved.

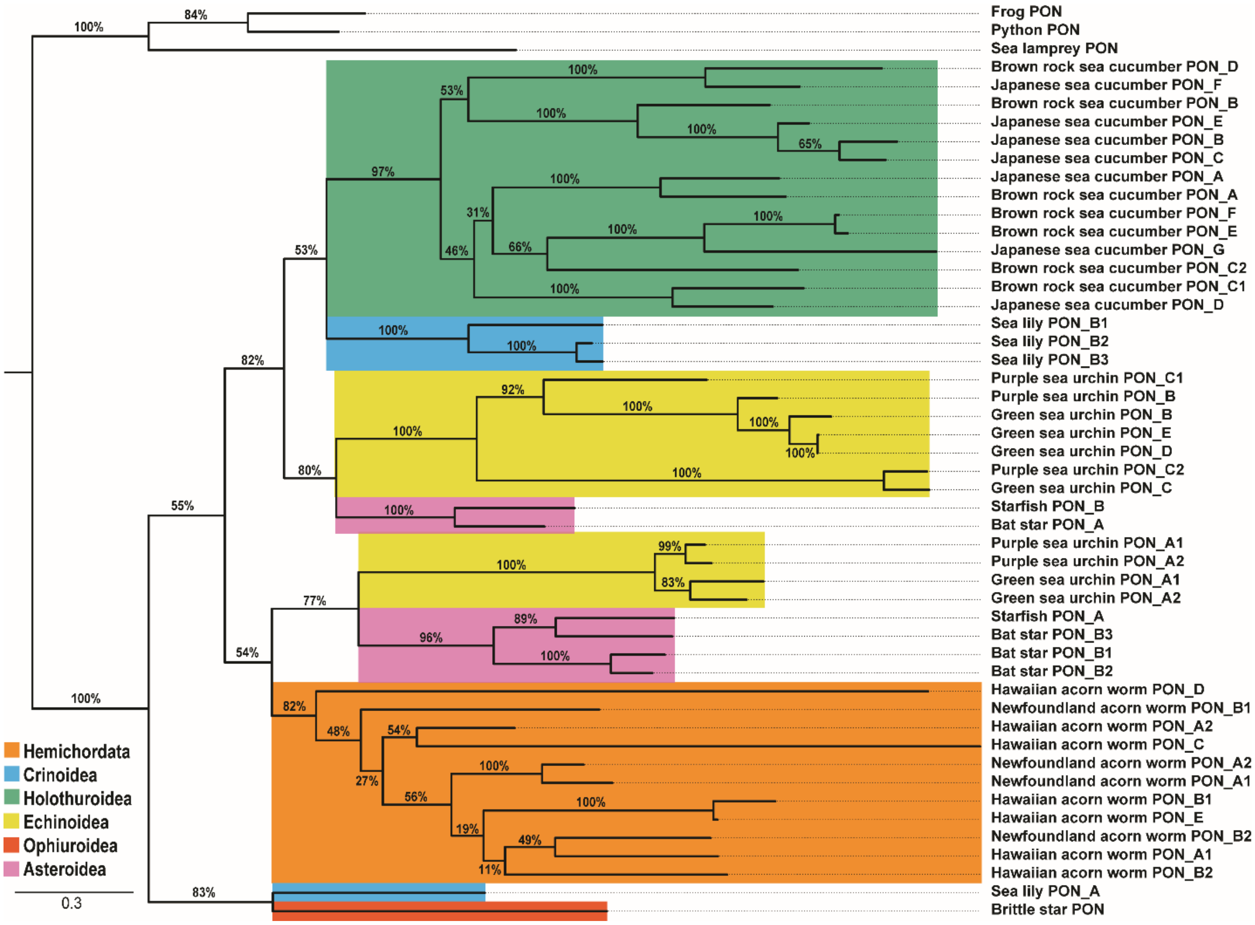


Supplementary Figure 4 Evolution of PON in ambulacraria with branch lengths. Phylogenetic tree of PON family proteins in ambulacraria determined by PhyML based on multiple sequence alignment. Bootstrap support values are shown as percentages out of 200, and the branch lengths have been scaled to reflect the relative branch lengths between all the sequences.
